# Transformation of acoustic information to sensory decision variables in the parietal cortex

**DOI:** 10.1101/2022.07.05.498869

**Authors:** Justin D. Yao, Klavdia O. Zemlianova, David L. Hocker, Cristina Savin, Christine M. Constantinople, SueYeon Chung, Dan H. Sanes

## Abstract

The process by which sensory evidence contributes to perceptual choices requires an understanding of its transformation into decision variables. Here, we address this issue by evaluating the neural representation of acoustic information in auditory cortex-recipient parietal cortex while gerbils either performed an auditory discrimination task or while they passively listened to identical acoustic stimuli. During task performance, decoding performance of simultaneously recorded parietal neurons reflected psychometric sensitivity. In contrast, decoding performance during passive listening was significantly reduced. Principal component and geometric analyses each revealed the emergence of decision-relevant, linearly separable manifolds, but only during task engagement. Finally, using a clustering analysis, we found subpopulations of neurons that may reflect the encoding of separate segments during task performance: stimulus integration and motor preparation or execution. Taken together, our findings demonstrate how parietal cortex neurons integrate and transform encoded auditory information to guide sound-driven perceptual decisions.

## Introduction

Integrating sensory information over time is one of the fundamental attributes that is required for accurate perceptual decisions (Brody and Hanks, 2016; Shadlen and Kiani, 2013). This process is supported by the transformation of stimulus representations into decision variables. In the case of auditory stimuli, prior to the formation of decision variables, the central representations of acoustic cues are gradually reconfigured along the auditory neuraxis. Thus, auditory neurons become more selective to contextually-relevant acoustic features as one ascends the central pathway into the auditory cortex (Wang, 2018). Ultimately, individual acoustic components merge into auditory objects to guide perception (Bizley and Cohen, 2013). Similarly, primary visual cortex neurons are selective to the stimulus orientation (Hubel and Wiesel, 1962, 1968), whereas higher cortices are selective for more complex characteristics (Rust and Dicarlo, 2010; DiCarlo et al., 2012; Movshon and Simoncelli, 2014). A hierarchical progression of sensory information processing is also seen across the somatosensory ascending pathway where receptive fields grow more complex (Iwamura, 1998). These hierarchically transformed neural signals are ultimately decoded downstream of sensory cortices for stimulus-dependent decisions (Gold and Shadlen, 2007; Bizley and Cohen, 2013; Tsunada et al., 2016; Gold and Stocker, 2017; Runyan et al., 2017; Town et al., 2018).

Studies in both non-human primates and rodents suggest that the parietal cortex integrates sensory inputs and transforms them into decision signals (Shadlen and Newsome, 2001; Freedman and Assad, 2006; Harvey et al., 2012; Runyan et al.,2017; Driscoll et al., 2017). The parietal cortex receives direct projections from primary or secondary sensory cortices (Hackett et al., 2014; Wilber et al., 2015), has been causally implicated in the performance of perceptual decision-making tasks (Hanks et al., 2006; Katz et al., 2016; Licata et al., 2017; Yao et al., 2020), and its activity typically reflects action selection (Andersen and Cui, 2009; Hwang et al., 2017). Furthermore, parietal neurons gradually increase their spiking activity over time epochs that scale with the accumulation of sensory evidence (Roitman and Shadlen, 2002; Gold and Shadlen, 2007; Kiani et al., 2008; Hanks et al., 2015; Zhou and Freedman, 2019). Thus, while parietal cortex activity reflects decision variables, the manner in which relevant sensory stimuli are represented prior to this transformation remains uncertain.

To dissociate encoding of stimuli from encoding of choice, we recorded neural activity from parietal cortex while gerbils performed an auditory discrimination task (Yao et al., 2020), and again during passive listening sessions, using the same acoustic stimuli in the absence of behavioral choice. During task performance, decoded parietal cortex population activity correlated with psychometric sensitivity. Furthermore, neural trajectories from parietal cortex neural responses revealed the temporal progression of low-dimensional encoding of acoustic information that transitioned to encoding of behavioral choices. During passive listening sessions, decoded parietal cortex population activity was poorer than decoding during task performance, but scaled with stimulus duration. The neural trajectories differentiated between each stimulus condition, but did not reflect a decision variable. Thus, the parietal cortex could accumulate auditory evidence for the purpose of forming a decision variable during task performance. Finally, our clustering analysis based on neuronal response properties revealed subpopulations of parietal neurons that may reflect separate segments during task performance: stimulus integration and motor preparation or execution. We propose that the parietal cortex integrates and transforms bottom-up sensory information into decision variables during task performance.

## Results

We trained gerbils (n = 5) to perform a single-interval, two-alternative forced-choice AM rate discrimination task (Yao et al., 2020). Gerbils were trained to self-initiate each trial by placing their nose in a cylindrical port for a minimum of 100 msec. On each trial, a 4- or 10-Hz AM signal was presented from an overhead speaker and animals approached the left or right food tray on the opposite side of the test cage. A food pellet reward was delivered when animals approached the left food tray following a 4-Hz AM signal or the right food tray following a 10-Hz AM signal (Figure 1A). To measure the minimum time necessary to accurately perform the task, AM stimulus duration was varied across trials (100-2000 msec), and performance was quantified as the proportion of correct trials. Figure 1B shows example psychometric functions from two different animals. In both examples, task performance improved with increasing stimulus duration and reached an optimum at ≥800 msec. Figure S1 displays psychometric functions for all 5 gerbils (n = 44 sessions). Minimum integration time was defined as the shortest stimulus duration at which animals discriminated between the two AM signals at a performance level of 0.76, which is equivalent to the signal detection metric, *d’* of 1 (Hacker and Ratcliff, 1979). The distribution of minimum integration times across all sessions is shown at the bottom of Figure S1. There was no significant difference between minimum integration times for the 4- and 10-Hz AM signals (Wilcoxon signed-rank test, p = 0.67; 4-Hz minimum integration time median: 391 msec; 10-Hz minimum integration time median: 402 msec), demonstrating animals were not biased to either stimuli.

**Figure 1.**
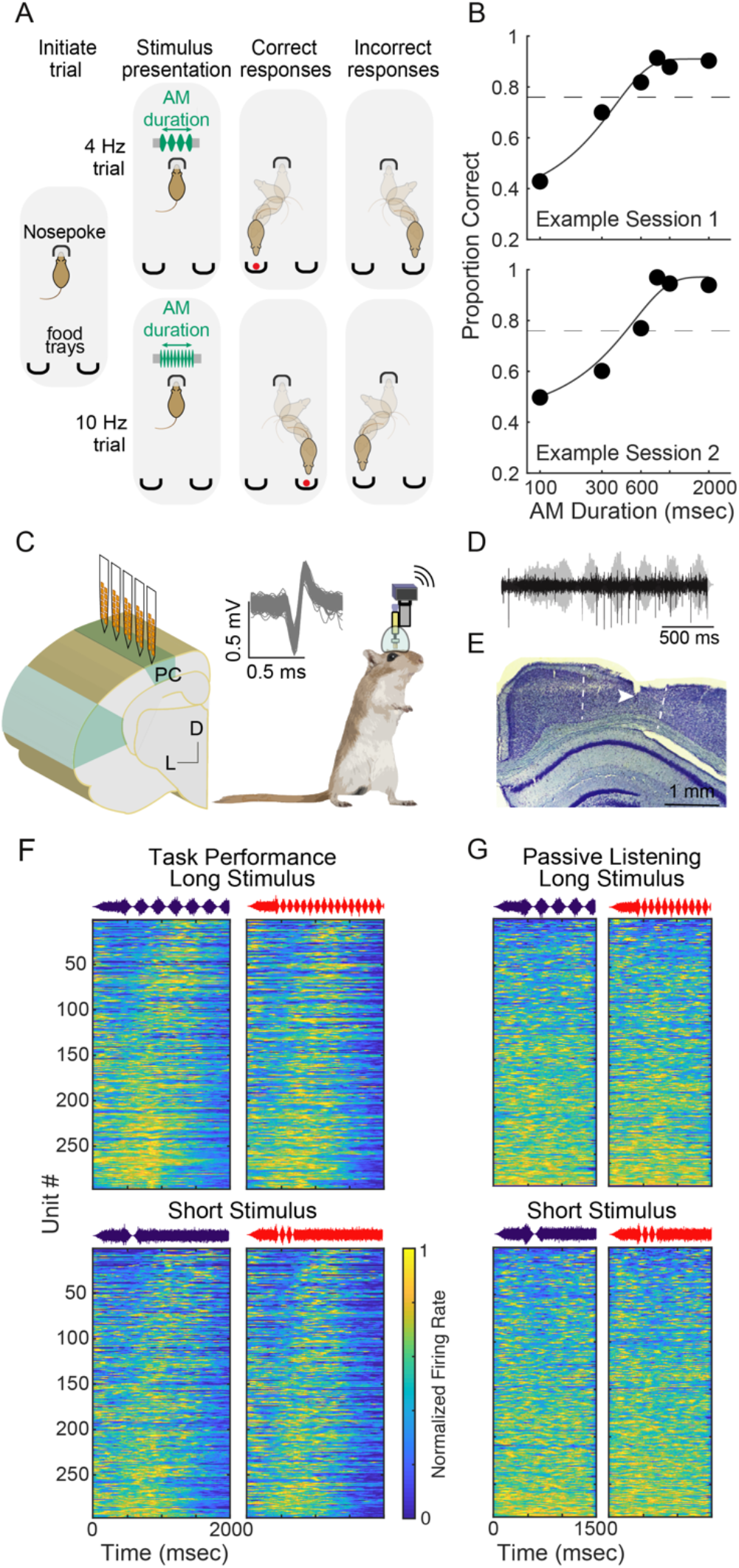
Behavioral measures of auditory task performance and neural recordings. **(A)** Schematic of the single-interval, two-alternative forced-choice AM rate discrimination task. Gerbils are required to discriminate between amplitude-modulated broadband noise presented at 4-versus 10-Hz across a range of stimulus durations (100-2000 msec). **(B)** Two example psychometric functions from two animals. **(C)** Chronic 64-channel electrode arrays were implanted into the left parietal cortex of 5 gerbils. **(D)** Raw waveform trace of neural response to AM signal. **(E)** Anatomically confirmed electrode track within the parietal cortex. **(F)** Normalized firing rate activity of all parietal cortex neurons during task performance sessions (n = 297) for “Long Stimulus” duration of 2000 msec (top) and “Short Stimulus” duration of 300 msec (bottom), sorted by time of maximum activity. **(G)** Same format as panel **F**. Normalized firing rate activity of parietal cortex neurons during the passive listening session (n = 284). Note that the total stimulus time is shorter for passive listening because trials did not exceed a total stimulus time of 1500 msec.

To assess parietal cortex neuron responses during task performance, we implanted trained animals with 64 channel electrode arrays, and obtained wireless recordings during auditory discrimination task performance. These recordings were compared to responses from the same neurons while animals listened passively to the identical acoustic stimuli (Figure 1C). Recorded physiological data (Figure 1D) were preprocessed to extract candidate waveforms for offline spike sorting procedures. One anatomically confirmed electrode track within the parietal cortex is shown in Figure 1E. We recorded a total of 297 units (22.9%, 68/297 classified as single-units) during task performance sessions and 284 units during passive listening sessions (22.9%, 65/284 classified as single-units). Figure S2 shows example post-stimulus time histograms (PSTHs) for one unit during one task performance and one passive listening session. The responses of all recorded units are shown in Figure 1F and G. Although there was a diversity of PSTH patterns during task performance, a fraction of parietal units displayed an initial decline in spike rate, followed by a gradual increase during the AM target stimuli. This temporal pattern of neural response was similar to that observed by parietal neurons during visual decision-making that also displayed ramping activity with increasing sensory evidence (Huk and Shadlen, 2005; Kiani et al., 2008; Okazawa et al., 2021). In contrast, a differential firing rate across time was not evident during passive listening sessions.

To determine whether parietal cortex population activity was sufficient to account for sound-driven task performance, we constructed linear classifiers using support vector machines (SVM), as described in Yao and Sanes (2018; see Methods). Briefly, AM discrimination was quantified across our parietal cortex population with a linear population readout scheme. The population linear classifiers were trained to decode responses from a proportion of trials to each individual AM rate signal (4-versus 10-Hz) across each stimulus duration (Figure 2A). Cross-validated classification performance was determined as the proportion of correctly classified held-out trials. This population decoder analysis was applied to our dataset in two ways. First, decoder performance was assessed from simultaneously recorded single- and multi-units within each behavioral session (i.e., “within-session” analysis; Figure 2B). Second, we assessed decoder performance for all units pooled across all behavioral sessions (Figure 2F).

**Figure 2.**
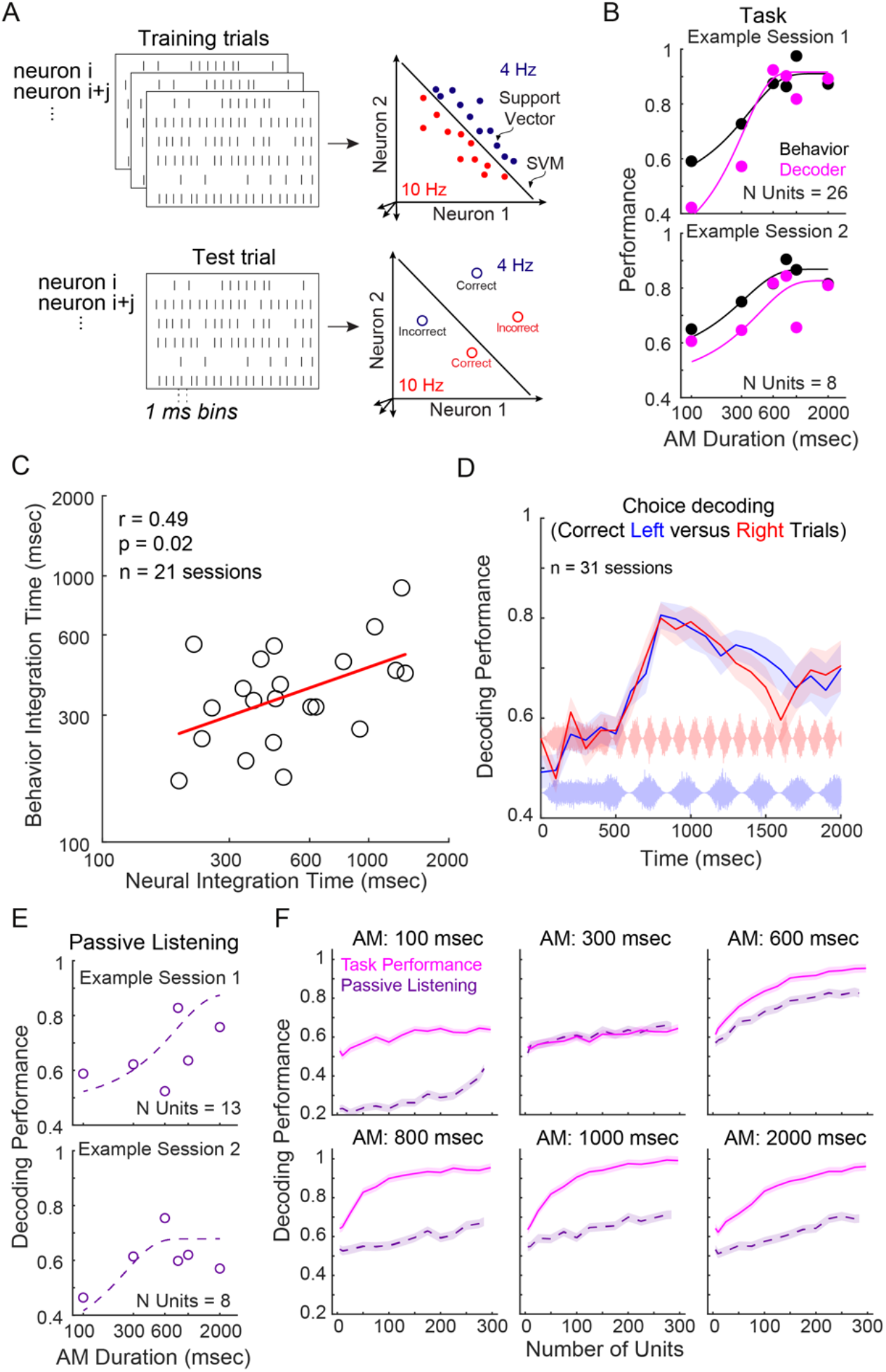
Parietal cortex population activity reflects auditory task performance and contains auditory information. **(A)** Schematic of linear population readout procedure. Population linear classifiers were trained to decode the responses from a subpopulation of simultaneously-recorded parietal cortex neurons from a proportion of trials to each AM rate signal (4 Hz versus 10 Hz) across each stimulus duration. Cross-validated classification performance was determined as the proportion of correctly classified held-out trials that were not used during classifier training. This procedure was performed across 250 iterations with new randomly drawn sampled train and held-out trials for each iteration. **(B)** Within-session population decoder results (pink) and corresponding behavior performance (black) from two example sessions from two animals during task performance. **(C)** Behavior versus neural integration times. Solid red line represents the linear regression. Pearson’s r and statistical significance are noted in the top-left corner of the figure panel. **(D)** Average ± SE within-session decoding performance for correct Left versus Right trials. **(E)** Within-session population decoder results from two example sessions from two animals during passive listening sessions. **(F)** Population decoding performance for task performance (pink) and passive listening (purple) conditions across increasing number of recorded units for each stimulus condition. A resampling procedure was used to randomly select a subpopulation of units with increasing increments of 25 across 500 iterations prior to applying the decoding readout procedure. Decoding performance for both session types increased with the number of units. For all stimulus durations except 300 msec, decoding performance during task performance exceeded that of passive listening sessions across all unit totals. For passive listening sessions, maximum decoding performance was found when including all recorded units, whereas decoding performance during task performance reached its peak when including ~≥80% of total units. Maximum decoding performance for task performance sessions asymptotes higher than passive listening sessions, but are comparable between 300-600 msec stimulus duration (Figure S3I), which is near the average behavioral integration time.

For the within-session analysis, we implemented a standard criterion to only assess sessions with a minimum of 5 simultaneously recorded single- and/or multi-units (n = 28/44 sessions). Figure 2B shows example within-session population decoder results from two animals during task performance. In both cases, decoding performance increases with longer stimulus durations, in line with psychometric performance. Decoding performance and corresponding behavioral performance for each stimulus type (4- and 10-Hz AM rate) across all sessions are shown in Figure 3SA. Neural minimum integration times were calculated as the stimulus duration corresponding to decoding performance of 0.76. The distributions of neural minimum integration times are plotted in Figure 3SB. We found a significant positive correlation between behavior integration times and corresponding decoder integration times (Figure 2C; r = 0.49, p = 0.02), and similar trends were observed for both trial types (Figure 3SC; 4-Hz AM: r = 0.54, p = 0.01; 10-Hz AM: r = 0.44, p = 0.05). This suggests parietal cortex activity reflects auditory-based decisions.

**Figure 3.**
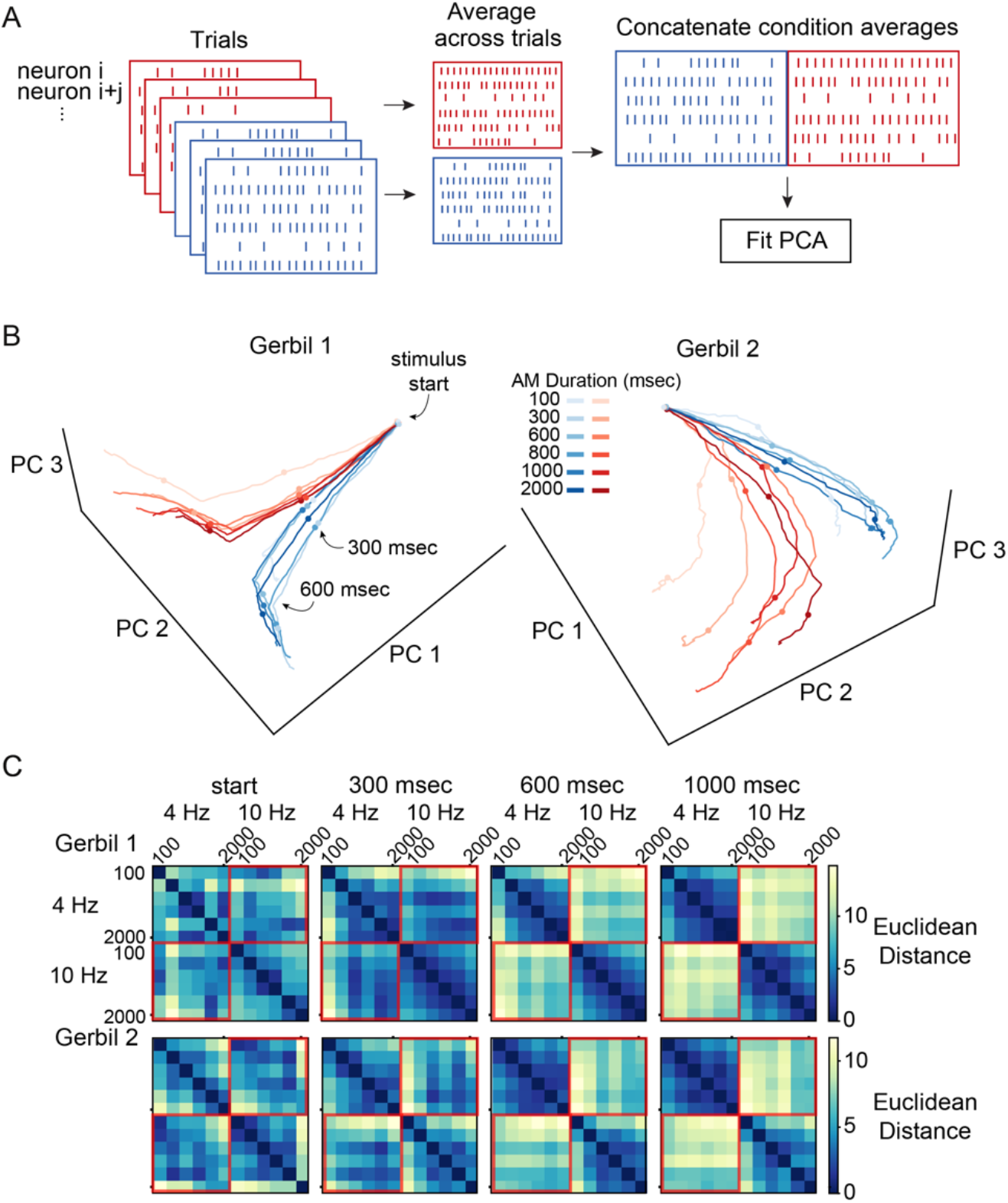
Parietal cortex population dynamics during task performance. **(A)** Principal component analysis (PCA) was performed on trial averaged neural responses from two of the five implanted gerbils. **(B)** Population activity plotted in a three-dimensional (3D) principal component space that originated from PSTHs of all recorded units for two gerbils during task performance. The neural trajectories (i.e., neural manifolds) in this state space correspond to the population responses across different times for separate AM rates and stimulus durations. Dot symbols represent 0, 300, and 600 msec after stimulus (AM) onset. **(C)** Distances between each stimulus condition, calculated using euclidean distance in the space spanned by the top 3 principal components, across time points of 0, 300, 600, and 1000 msec after AM onset.

We next asked whether the time course of decoder performance aligned with behavioral integration times. Figure 2D displays average ± SEM decoding performance as a function of time for correct left versus right trials across all 31 recorded sessions. At trial onset, decoding performance was near chance, and increased to a peak value of ~0.80 (average ± SEM, Left: 0.81 ± 0.03; Right: 0.80 ± 0.03). The maximum decoding performance occurred at ~350 msec of AM signal duration which is nearly identical to the average behavioral integration time (378 msec; Figure S1). To illustrate choice-related activity across trial durations, we also plotted decoding performance as a function of time relative to response latency (Figure 3SD). Decoding performance gradually begins to increase ~1000 msec prior to response latency and decoding performance peaks ~600 msec prior to response latency.

Although the striking alignment of neural and behavioral performance suggests that auditory information is being integrated within the parietal cortex, it does not provide a direct measure of stimulus coding. Therefore, we recorded from the same parietal neurons studied during task performance while animals listened passively to the identical 4 and 10 Hz AM stimuli (Figures 1G and S2B). Within-session population decoder results for two passive listening sessions are shown in Figure 2E. Decoding performance across all sessions for each stimulus type are shown in Figure S3E. The distributions of neural minimum integration times during passive listening sessions are plotted in Figure S3F. Overall, only a fraction of passive listening sessions yielded minimum integration times (n = 17/29; maximum decoding performance did not reach 0.76 for the remaining 12 sessions). For these passive listening sessions, decoding performance scaled with increasing stimulus duration (Figure 2E and S3E), suggesting that the parietal cortex could accumulate this sensory evidence for the purpose of forming a decision variable.

To directly compare decoder performance during task performance and passive listening, we examined the 19 instances where both types of session fit the criterion of 5 simultaneously recorded units. In the majority of those instances (13/19 sessions), minimum integration time was better or could only be calculated during task performance (Figure S3G). In 6 cases, integration time diminished or could not be calculated during task performance (Figure S3H). This suggests that the parietal cortex is more strongly engaged when animals are required to integrate sensory information for decision-making.

We further examined decoding performance as a function of the number of total recorded units for task performance and passive listening sessions (Figure 2F). A subsampling procedure was applied to randomly select a subpopulation of units (25-297 for task performance and 25-284 for passive listening sessions; increasing increments of 25) across 500 iterations. During each iteration of the resampling procedure, a new subpopulation of units was randomly selected (without replacement) prior to the decoding readout procedure. For each stimulus duration, decoding performance for both task performance and passive listening session types increased with the number of units, demonstrating evidence for population-level encoding.

If the parietal cortex does compute a decision variable, then its population activity should gradually evolve into two independent patterns. To test this idea, we assessed the dynamics of parietal cortex population activity by applying principal components analysis to the trial averaged neural responses (Figure 3A). This analysis was conducted on two of the five recorded animals as they both provided the large majority of recorded units (Gerbil 1 n = 115/297 units; Gerbil 2 n = 165/297 units). Figure 3B depicts population activity in a three-dimensional (3D) principal component space that originated from PSTHs of recorded units from two animals during task performance (top 3 principal components, explained variance: Gerbil 1 = 79.7%; Gerbil 2 = 88.8%). The neural trajectories in this state space correspond to the population responses across different times for each AM rate and the stimulus durations. At stimulus onset, neural trajectories started at a similar position, but began to diverge toward the relevant decision subspace (4 Hz versus 10 Hz) after ~300 msec of acoustic stimulation. This divergence toward the relevant decision subspace over time is further demonstrated when we measured the euclidean distance between each pair of trajectories in the space spanned by the top three principal components (Figure 3C). Over time, the distance between the trajectories that correspond to the stimulus durations of opposing AM rates (4 Hz versus 10 Hz) dramatically increased (Figure 3C, upper right and lower left quadrants of each matrix; outlined in red), while the average distance between the trajectories that correspond to the stimulus durations within each AM rate remained low (Figure 3C, upper left and lower right quadrants of each matrix). In other words, the resulting distance matrix is block diagonal showing that trajectories corresponding to the same choice remained closer to each other than to the trajectories corresponding to the opposite choice – thereby indicating the existence of a choice manifold. We define “manifold” as the collection of population neural responses that encode a stimulus. These results are consistent with the integration of sensory evidence over time and the representation of a decision variable (DV) by the neural population.

In principle, decision variables should not be computed when animals are disengaged from a sensory task. To test this idea, we performed the same PCA analysis to trial averaged neural responses recorded during passive listening sessions. Figure 4A depicts population activity in 3D principal component space that originated from PSTHs of recorded units from two animals during passive listening (top 3 principal components, explained variance: Gerbil 1 = 82.3%; Gerbil 2 = 85.2%). In contrast to the neural trajectories of parietal cortex neural responses during task performance, the neural trajectories elicited during passive listening did not diverge according to the two separate manifolds corresponding to AM rates (Figure 3C versus Figure 4B, outlined red squares). This is reflected in the difference in decoding performance between the two behavioral conditions (Figure 2B, E, and F). Instead, the neural trajectories elicited by each combination of AM rate and stimulus duration eventually occupied separate positions in the principal component space. This is further demonstrated by the differences in distances between each stimulus condition (Figure 4B). Combined with the finding that several decoding sessions yielded an integration time (n = 17/29; Figure S3F), this suggests that, while acoustic information is encoded in the parietal cortex during passive listening, decision variables are not computed.

**Figure 4.**
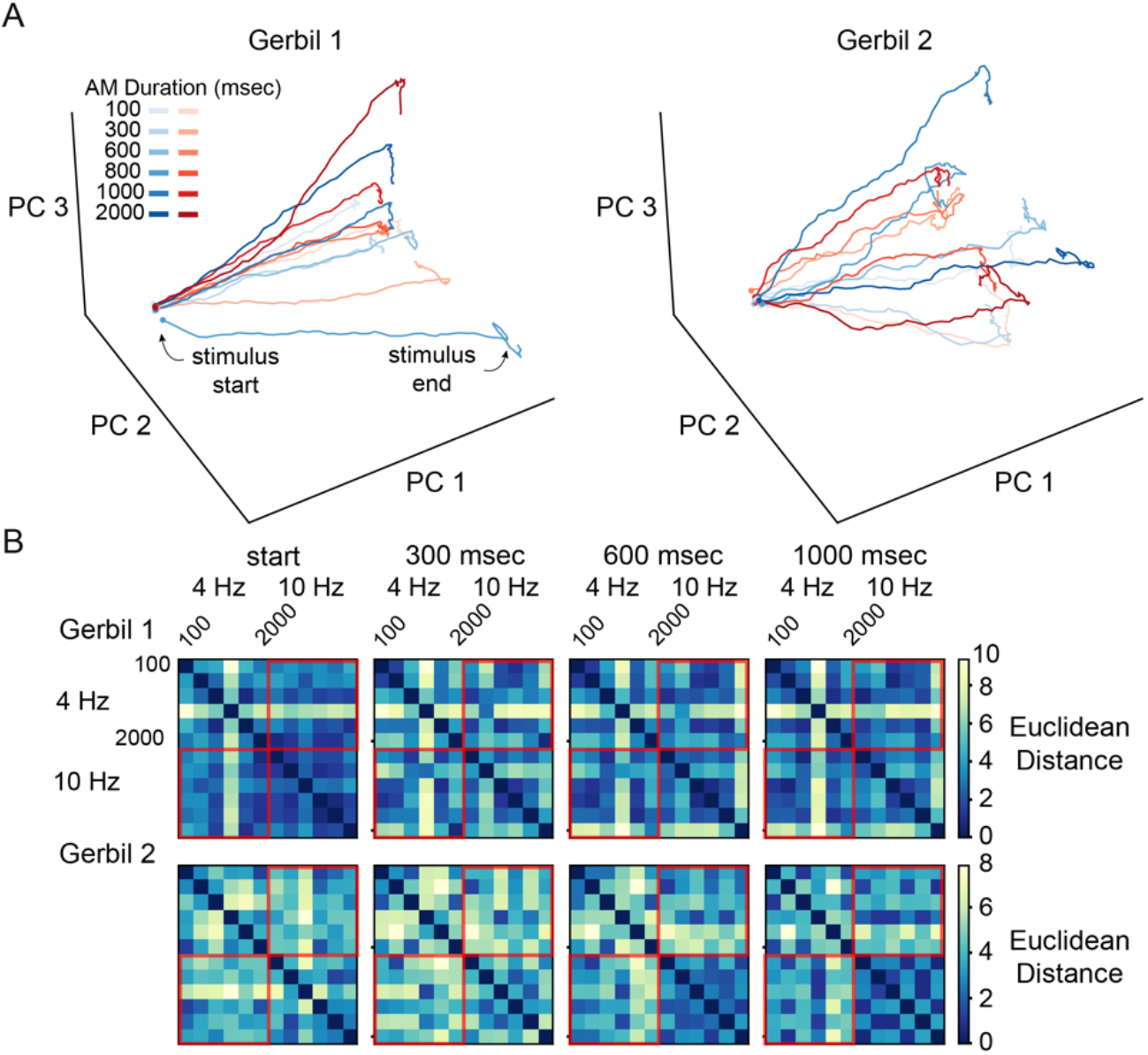
Parietal cortex population dynamics during passive listening. **(A)** Population activity plotted in a three-dimensional (3D) principal component space that originated from PSTHs of all recorded units for two gerbils during passive listening. The neural trajectories (i.e., neural manifolds) in this state space correspond to the population responses across different times for separate AM rates and stimulus durations. **(B)** Distances between each stimulus condition, calculated using euclidean distance in the space spanned by the top 3 principal components, across time points of 0, 300, 600, and 1000 msec after AM onset.

Previous studies have hypothesized that the brain transforms sensory information into linearly separable representations (Cohen at al., 2020; Chung and Abbott, 2021). This “untangling” of representations has been suggested to be a more prominent feature of higher-order brain areas (Cohen at al., 2020; DiCarlo and Cox, 2007). Thus, the segregation of neural trajectories from parietal cortex activity into two separate subspaces during task performance may represent an encoding strategy that enables linear readout of decision variables (Figure 5A). To test whether the neural representations of the AM rate stimuli in principal component space are consistent with this prediction, we employed three measures of “untangling”: capacity, manifold radius and manifold dimensionality (Chung et al., 2018). These three measures define the separability of objects based on their neural manifolds. Capacity measures how many different object classes can be linearly separated with high probability. Manifold radius and dimensionality quantify the size of the manifold; essentially, the variance of the points that belong to the manifold as well as its spread along different axes. To compute these measures, we first binned the spiking activity into 80 msec bins and then split the trial activity into 2 conditions according to the AM rates, 4- and 10-Hz, combining across stimulus durations as well as all of the animals. Consistent with the idea of untangling, we found that there was an increase in capacity (Figure 5B) and a decrease in manifold radius and dimensionality after the onset of the AM stimulus (Figure 5C and D). Finally, we computed the norm of the manifold center over time, which measures the distance of the center of the manifold to the origin, to understand if the object manifolds corresponding to each stimulus frequency move over time. This measure increased (Figure 5E), suggesting that the 4- and 10-Hz manifolds moved away from their starting position over time. These trends continue until about 600 msec into the AM stimulus (1000 msec of absolute trial time) at which point all four measures plateau. Importantly, the time course of this change in neural representation is consistent with the time window over which the animal has to accumulate evidence and make its decision. Together, our findings suggest that the transformation of sensory evidence into decision variables in the parietal cortex is accompanied by changes in the neural representation that supports the separability of the stimulus manifolds.

**Figure 5.**
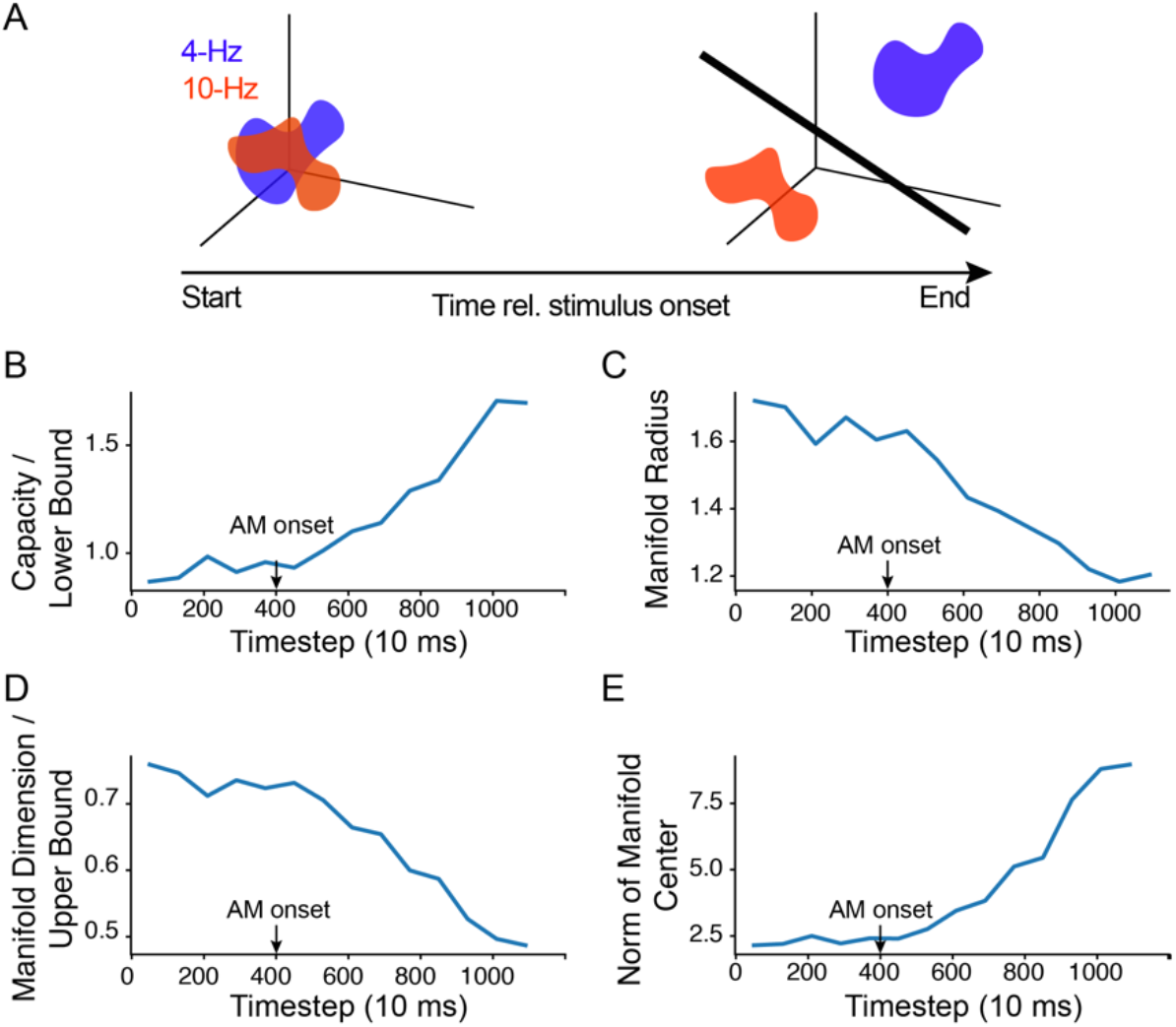
“Untangling” of neural responses. **(A)** 4- and 10-Hz stimulus manifolds (blue and red clouds) are entangled at the beginning of the trial but become more linearly separable over the course of the stimulus integration window. **(B)** Change in the mean manifold capacity over time. Amplitude modulated sound onset is at 400 msec. Increasing capacity corresponds to an increase in linear separability of the manifolds. **(C)** Change in the mean manifold radius over time. Decreasing radius corresponds to an increase in linear separability of the manifolds. **(D)** Change in the mean manifold dimensionality over time. Decreasing manifold dimensionality corresponds to an increase in linear separability. **(E)** Change in the mean norm of manifold center over time. Increasing norm of manifold center corresponds to manifolds moving further away from the origin.

Up until this point, we have assumed that the recorded population was homogeneous. Therefore, we asked whether there were distinct functional classes and, if so, whether they differentially represented the decision variable. To test this, we performed clustering on PSTHs (Raposo et al., 2014; Namboodiri et al., 2019; Hocker et al., 2021). Specifically, for each neuron we averaged over trials with the same AM stimulus rate (4- or 10-Hz) to obtain two conditional PSTHs spanning the 2 seconds after trial initiation. We then concatenated these 2 PSTHs to create a high-dimensional feature space that represents the unique activity of each neuron across the two stimulus conditions. Analysis of the angles between these data points in feature space indicated that there were clusters in this dataset (PAIRS test, Raposo et al., 2014), and further analysis using the gap statistic revealed 3 subpopulations of neurons in the population response (Figure 6A-D). Cluster 1, the largest cluster in the population, demonstrated activity at the onset of unmodulated noise (i.e., the 400 ms before the AM stimulus), and persisted through the trial (Figure 6A, B). Clusters 2 and 3 displayed decreased activity at the beginning of a trial, with a ramping of activity peaking at ~1s, which is approximately the time that animals had already made their initial movement towards the reward ports. Clusters 1 and 3 were well-represented, though Cluster 2 was predominantly represented in only one gerbil. We confirmed that this clustering result is robust to different forms of pre-processing of the responses, such as downsampling to coarse time bins, or smoothing over multiple time bins (Figure S4A-D). Clustering on the passive condition reveals the presence of 2 clusters, though the population predominantly belonged to only a single cluster (Figure S5).

**Figure 6.**
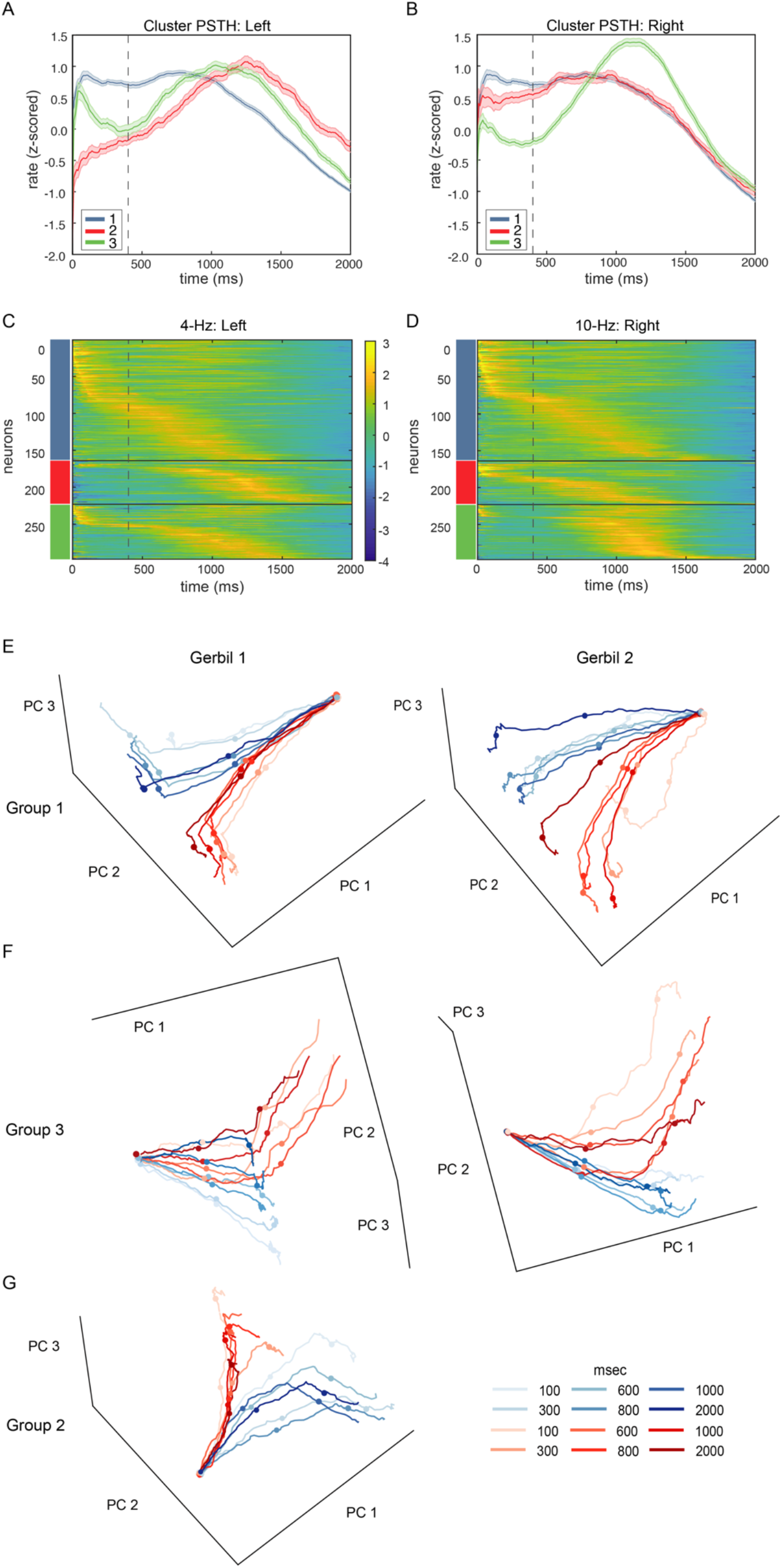
Clustering of conditional PSTH responses reveals three distinct subpopulations of neurons in the parietal cortex. **(A)** Cluster-averaged PSTH for 4-Hz AM stimulus indicating a reward at the left reward port. **(B)** Similar to (**A)**. 10-Hz AM stimulus indicating food at the right reward port. Dotted line indicates onset of the 4- and 10-Hz AM stimulus, and error bars denote s.e.m. **(C)** Population PSTH responses for 4-Hz AM stimulus. **(D)** Similar to **(C)** except for 10-Hz AM stimulus. Responses are grouped by cluster identity (colored rectangles), and sorted by time-to-peak within each cluster, for each stimulus condition. **(E)** Population activity plotted in a 3D principal component space that originated from PSTHs of “Group 1” units for two gerbils during task performance. **(F)** Population activity plotted in a 3D principal component space that originated from PSTHs of “Group 3” units for two gerbils during task performance. **(G)** Population activity plotted in a 3D principal component space that originated from PSTHs of “Group 2” units for one gerbil during task performance. Dot symbols represent 0, 300, and 600 msec after stimulus (AM) onset.

To evaluate the computational roles of each neural cluster and determine whether specific subpopulations of neurons reflect the integration of sensory evidence and the formation of decision variables, we performed principal component analysis on each cluster individually. Specifically, the principal component analysis was fit to trial averaged neural responses across time for the 3 clusters separately for 2 of the 5 gerbils. The neural trajectories in this state space correspond to the population responses for each cluster across different times for separate AM rates and stimulus durations. We show the neural trajectories for each cluster in the space spanned by the top 3 principal components of each respective cluster (Figure 6E-G). The neural trajectories across all clusters show a separation between the two AM rates, but diverge at separate time points. This suggests that each cluster may encode different task-relevant information. For example, the neural trajectories in the principal component space of Clusters 1 (Figure 6E) and 2 (Figure 6G) display strong divergence between the corresponding AM rates much sooner than the neural trajectories of Cluster 3 (Figure 6F). Specifically, the divergence of neural trajectories for Clusters 1 and 2 occurs within 600 msec after stimulus onset, which corresponds to the behaviorally-relevant integration times (Figure S1). This suggests that Clusters 1 and 2 may reflect the transformed sensory signals received from the auditory cortex during task performance. The neural trajectories for Cluster 3 diverge later than ~600 msec after stimulus onset (>~1000 msec total trial duration), suggesting that Cluster 3 may reflect the motor preparatory signal that is executed during the task. Overall, our results demonstrate distinct subpopulations of parietal cortex neurons encode separate segments of task performance: stimulus integration and motor preparation or execution.

## Discussion

Our central finding is that, during task performance, parietal cortex neurons integrate and transform behaviorally-relevant acoustic information to drive sound-driven perceptual choices. Decoded parietal cortex activity reflected psychometric sensitivity during task performance and aligned with behavioral measures of integration time. In contrast, decoded neural activity from passive listening sessions was dramatically reduced (Figure 2F). To analyze whether parietal cortex activity could support sensory evidence accumulation, we applied principal component and geometric analyses, and found the emergence of decision-relevant, linearly separable manifolds that reflect behavioral integration time during task performance. Taken together with our previous finding that auditory cortex projections to the parietal cortex play a causal role in producing behavioral integration time (Yao et al., 2020), we propose that the parietal cortex transforms auditory afferent input into decision variables.

Previous work shows that parietal cortex neurons are strongly modulated by the behavioral relevance and context of acoustic stimuli (Stricanne et al., 1996; Nakamura, 1999; Grunewald et al., 1999; Linden et al., 1999). Furthermore, functional interactions of simultaneously recorded parietal neurons are greater than that seen among auditory cortex neurons and also extend to longer time scales (Runyan et al., 2017), demonstrating the transition to somewhat more behaviorally-relevant timescales for processing sensory information into stimulus-driven decisions. These functional properties are clearly associated with anatomical connectivity between primary or secondary auditory cortices to parietal cortex, which are strongly apparent across many species (Pandya and Kuypers, 1969; Reep et al., 1994; Rauschecker and Scott, 2009; Wilber, et al., 2014; Song, et al., 2017; Yao et al., 2020).

Our current results complement these findings by demonstrating how auditory encoded information is transformed from an uninformative representation during passive listening, to meaningful integration times that reflect behavioral performance. During passive listening, decoded activity from simultaneously recorded parietal cortex neurons is poorer than decoded activity during task performance, but scales with stimulus duration (Figure 2E). Thus, evidence for encoded sensory inputs within parietal cortex derives from the scaling of decoded parietal cortex activity with the amount of presented stimulus information. Our principal component analyses further demonstrate an even greater difference of parietal cortex activity between behavioral conditions. Whereas encoded auditory information from parietal neurons occupied separate positions in subspace during passive listening (Figure 4A, B), we found an emergence of decision-relevant, linearly separable manifolds on a behaviorally-relevant timescale during task performance (Figure 3B, C). This is specifically demonstrated by a clear separation of manifolds that correspond to the two AM rates (4- and 10- Hz), which is also reflected by two separate decision outcomes (e.g., approach the left or right food tray). The transition of sensory encoding between passive and task-engaged contexts suggests that sensory information transitions into a decision-making context and reflects the learned association between sensory categorization and motor execution. This is in contrast to categorical sensory representations (Banno et al., 2020), which would be true if parietal cortex neurons represented stimulus categories during passive listening conditions. It is worth noting that the recordings for the passive condition were collected from the same highly trained animals, so the differences in representation cannot be explained by the lack of association between stimulus and choice.

Our principal component and geometric analyses demonstrated that decision variables emerge within parietal cortex activity during task performance. We predict that the role of the parietal cortex is to transform stimulus information into a representation that can be easily decoded into action. While the neural manifolds that correspond to 4- and 10-Hz AM rates increasingly diverge during task performance (Figure 3B), these representations become more “untangled,” or linearly separable, over behaviorally-relevant timescales (Figure 5A-E). This is computationally desirable because it suggests that the parietal cortex can read out auditory information using the simplest possible decoder. This result is consistent with predictions from artificial neural network models for auditory processing (Stephenson at al., 2019) and is also consistent with the previous findings that individual neurons show mixed selectivity for task variables (Okazawa et al., 2021) and may change activity patterns without affecting the overall ability of the population to encode the relevant task information (Driscoll et al., 2017). Finally, our result shows that this untangling of representations occurs not only by compression in dimensionality and size of the stimulus manifolds, but also by the stimulus manifolds moving away from each other.

Clustering on the temporal responses of parietal cortex neurons during task performance revealed 3 subpopulations of neurons, with two clusters being well-represented. One sub-population (“Cluster 1”) demonstrated the encoding of AM stimulus information, while the other dominant sub-population (“Cluster 3”) displayed a gradual increase in activity that peaked ~1 sec of total trial duration (~600 msec after stimulus onset), roughly around the time animals initiated their movement for reward retrieval. This late-in-trial segment is likely related to preparatory movement activity, and is distinctly separate from neurons that integrate stimulus information. A third cluster (“Cluster 2”) seemed to share a similar phenotype with Cluster 1 since its corresponding neural trajectories diverged relatively around the same time. However, when comparing PSTHs, Cluster 2 neuronal responses were more modulated for ipsilateral (left; 4-Hz) conditions, relative to Cluster 1. It is important to note that this cluster was only present in 1 gerbil. It is possible that neurons from Cluster 2 may belong to Cluster 1, or alternatively, observing ipsilateral encoding of sensory evidence integration is simply a rare type of response property in the parietal cortex.

Previous studies did not find separable clusters when examining the mixed selectivity of parietal cortex activity (Raposo et al., 2014). In that work, PAIRS analysis was performed on time-averaged activity across different sensory stimuli (auditory and visual), and found that responses were not separable into distinct subpopulations. Our results in parietal cortex responses instead used time-dependent response profiles that were restricted to a single stimulus modality to analyze potential clustering. We believe that these two results do not conflict, and that taken together they highlight how clustering is a flexible tool to characterize a variety of encoding properties across subpopulations of neurons.

While our study focused on the sensory input to the parietal cortex, it did not address the neural mechanism that causes the transformation of sensory representations into decision variables. This process is thought to require descending input from the prelimbic region of the frontal cortex (Wilber et al., 2015; Granon and Poucet, 2000). The cingulate cortex may provide one source of task-relevant information to the parietal cortex as neurons can encode context-dependent signals, which can be read out by locus coeruleus activity (Joshi and Gold, 2022). This provides a potential neural circuit for appropriately modulating parietal cortex activity during task performance where represented encoded sensory information is integrated, grouped, and transformed into decision variables that can be projected to motor circuits (Harvey et al., 2012).

Our results do not indicate whether the formation of sensory-driven decisions also occurs in premotor circuits, such as those that strictly involve action planning, including striatal circuits (Cox and Witten, 2019). Furthermore, our results do not demonstrate whether the auditory temporal integration signals observed are exclusively computed within parietal cortex, or reflect a computation performed elsewhere in the brain that are contingent on motor execution, such as brainstem networks (Horwitz and Newsome, 1999; 2001; Horwitz et al., 2004; Felsen and Mainen, 2008). Future work will determine whether the transformation of sensory integrated signals into task-engaged choice-specific variables occurs within separate neural circuits and/or is dependent on the execution of motor actions, such as the movements associated to report the decisions (Freedman and Assad, 2011).

In summary, the representation of behaviorally-relevant auditory information occurs in the parietal cortex even when animals are passively listening to the stimuli. However, it is only during task engagement that this information is transformed to a decision variable that correlates with psychometric performance. We demonstrated this with principal component and geometric analyses, each of which show that sensory evidence is accumulated across a time frame that matches behavioral integration time (Yao et al., 2020). Thus, our findings provide a plausible argument for the parietal cortex’s role in integrating and transforming encoded auditory information into decision variables to guide sound-driven behavior.

## Materials and Methods

Adult Mongolian gerbils (*Meriones unguiculatus,* n = 5, 3 males) were weaned from commercially obtained breeding pairs (Charles River). Animals were housed on a 12 h light/12 h dark cycle and provided with ad libitum food and water unless otherwise noted. All procedures were approved by the Institutional Animal Care and Use Committee at New York University.

### Behavior

#### Behavioral apparatus

Adult gerbils were placed in a plastic test cage (0.4 x 0.4 x 0.4 m) in a sound-attenuating booth (GretchKen Industries, Inc; internal dimensions: 1.5 x 1.5 x 2.2 m) and observed via a closed-circuit monitor. Acoustic stimuli were delivered from a calibrated free-field tweeter (DX25TG0504; Vifa) positioned 1 m above the test cage. Sound calibration measurements were made with a 1/4-inch free-field condenser recording microphone (Brüel & Kjaer) placed in the center of the cage. Stimulus, food reward delivery, and behavioral data acquisition were controlled by a personal computer through custom MATLAB scripts (written by Dr. Daniel Stolzberg: https://github.com/dstolz/epsych) and an RZ6 multifunction processor (Tucker-Davis Technologies).

Psychophysical training and testing was implemented with a positive reinforcement appetitive one-interval alternative forced-choice (AFC) procedure, as described previously (Yao et al., 2020). Briefly, gerbils were placed on controlled food access and trained to discriminate between amplitude modulated (AM) frozen broadband noise (25-dB roll-off at 3.5 kHz and 20 kHz) at 4-versus 10-Hz at 100% modulation depth. Each AM stimulus were presented at a sound pressure level (SPL) of 66 dB and had a 200 ms onset ramp, followed by an unmodulated period of 200 ms that transitioned to an AM signal for a set duration, followed by an unmodulated period. Gerbils self-initiated trials by placing their nose in a cylindrical port (nose poke) for a minimum of 100 ms that interrupted an infrared beam and triggered an acoustic stimulus. During acoustic stimulation, a gerbil approaches the left or right food tray and the infrared beam at the correct food tray is broken, a pellet dispenser (Med Associates) delivers one reward dustless precision pellet (20 mg; Bio-Serv). Gerbils were first trained to distinguish between 4-versus 10-Hz AM with a stimulus duration of 2,000 ms (proportion of trials correct > 0.85) across two sessions, and then were presented with shorter durations (e.g., 1,000, 800, 600, 300, and 100 ms) across subsequent sessions. For each trial, the probability of a 4- or 10-Hz AM stimulus presentation is 50% and its duration is a random draw. Figure 1A displays the schematic of the task.

During sessions to assess perceptual sensitivity, six signal durations for each of the 4- and 10-Hz AM stimuli (100, 300, 600, 800, 1,000, and 2,000 ms) are presented. Integration time is assessed by examining how performance scales with stimulus duration. Proportions of correct trials across stimulus durations for each AM rate are fitted with psychometric functions using the open-source package psignifit 4 for MATLAB (Schutt et al., 2016). Psychometric functions of the proportion of correct trials are plotted as a function of stimulus duration. Minimum integration time was defined as the stimulus duration at which proportion of correct trials = 0.76, which is equivalent to the signal detection metric, *d*’, equal to 1 (Hacker and Ratcliff, 1979).

### Electrophysiology

Extracellular single- and multiunit activity was recorded from the left medial parietal cortex. After gerbils were trained in the behavioral task, a silicon probe with 64 recording sites was implanted into the left medial parietal cortex (Neuronexus, model Buszaki64_5×12-H64LP_30mm). We targeted the medial portion of the parietal cortex because of its robust auditory cortex-recipient anterograde labeling (Yao et al., 2020). The probe was attached to a manual microdrive (Neuronexus, dDrive-XL) that allowed the electrode to be advanced and retracted. Probes were inserted at a 0- to 10-degree angle on a mediolateral axis. Typically, we aimed the rostral-most shanks of the array to be positioned at 3.3-3.6 mm rostral and 2.5 mm lateral to lambda. The surgical implantation procedure was performed under isoflurane anesthesia. Animals recovered for at least 1 week before being placed on controlled food access for further psychometric testing. At the termination of each experiment, animals were deeply anesthetized with sodium pentobarbital (150 mg/kg) and perfused with phosphate-buffered saline and 4% paraformaldehyde. Brains were extracted, post-fixed, sectioned on a vibratome (Leica), and stained for Nissl. Brightfield images were inspected under an upright microscope (Revolve Echo) and compared to a gerbil brain atlas (Radtke-Schuller et al., 2016) to verify targeted medial parietal cortex.

Physiological data were acquired telemetrically from freely-moving animals with a wireless headstage and received (W64, Triangle Biosystems). Analog signals were preamplified and digitized at a 24.414 kHz sampling rate (PZ5, Tucker-Davis Technologies) and fed via fiber optic link to the RZ5 base station (Tucker-Davis Technologies) and PC for storage and post-processing. Offline, electrophysiological signals underwent a common average referencing procedure (Ludwig et al., 2009) and bandpass filtered at 300-5000 Hz. Significant noisy portions of the signal that were induced by extreme head movements were removed by an artifact rejection procedure. An open source spike package (KiloSort; Pachitariu et al., 2016) was used to extract and cluster spike waveforms. Manual inspection of spike waveforms was conducted in Phy (Rossant et al., 2016). Well-isolated single units displayed clear separation in principal component space and possessed few refractory period violations (<10%). Units that did not meet these criteria were classified as multi-units. All sorted spiking data were analyzed with custom MATLAB scripts. Recordings were made both during task performance, and during passively listening sessions that occurred after task performance. All passively listening sessions were recorded immediately after each recorded task performance session with the nose poke and food trays removed from the test cage.

### Neural analyses

#### Population coding

We used a previously employed linear classifier readout procedure (Yao and Sanes, 2018) to assess AM rate discrimination across a population of parietal cortex neurons. Specifically, a linear classifier was trained to decode responses from a proportion of trials to each stimulus condition (e.g., 4 versus 10 Hz; Figure 2A). Spike count responses from all recorded neurons were counted within 100 msec time windows across the entire trial durations and formed the population “response vector”. Since the number of trials were typically unequal between stimulus conditions, we randomly subsampled (without replacement) a proportion of trials (i.e., 15 trials) from each unit. A support vector machine (SVM) procedure was used to fit a linear hyperplane to 80% of the data set (“training set”). Cross-validated classification performance was assessed on the remaining 20% across 250 iterations with a new randomly drawn sampled train and test sets for each iteration. Performance metrics were computed to determine the proportion of correctly classified and misclassified trials using an expanding time window (100 msec increments) across the entire trial duration. We restrained the analysis time window to correspond to each stimulus duration up to 600 msec (e.g., maximum time window did not exceed 300 msec for stimulus duration of 300 msec). A maximum time window of 600 msec was utilized for stimulus durations > 600 msec to control for movement-related signals that may arise when animals approach their selected food tray. This was particularly the case during task performance sessions. The SVM procedure was implemented in MATLAB using the “fitcsvm” and “predict” functions with the “KernelFunction” set to “linear”. This analysis was conducted for task sessions with ≥ 5 simultaneously recorded units and performance on easiest trial conditions (e.g., 2000 msec AM duration) ≥ 85% (n = 27/44 total sessions; median simultaneously recorded units = 6; interquartile range = 6.25).

#### Population response manifolds

Principal component analysis (PCA) was performed separately for both task performance and passive listening session types for each animal using trial averaged PSTHs. Trial averages were computed by binning the spiking data into 10 msec bins and using a rolling mean with a 50 msec window for each of 12 conditions (2 stimulus AM rates X 6 Durations) with each unit contributing to the PSTH for each condition. The PSTHs focused on the period of decision formation and spanned 400-1400 ms after stimulus onset (0-1000 msec after AM onset). For each unit, we concatenated the PSTH for each condition into a matrix of size N x CT where C is the number of conditions (2 AM rates x 6 durations), T is the number of time points (10 msec resolution) and N is the number of units. Each row of the matrix was then z-scored since PCA is known to be sensitive to overly active units. For the task performance condition, only trials where the animal made the correct choice were used in the analysis. Trials with only one spike in the time window of interest were left out from analysis. Units that had no data for one or more of the conditions were left out from the analysis. Finally, the analysis included all units recorded either simultaneously or separately. We confirmed that the same qualitative results (the presence of decision subspace during task performance and lack of one during passive listening) were obtained using simultaneously recorded units only.

#### Calculating distance between neural trajectories

To quantify the distance between neural trajectories (Figure 3C and 4B), we performed a bootstrapping procedure where the trial average was calculated over a random subsample of 10 trials per condition and this was done 1000 times to create 1000 new trial averages per condition per unit. These conditions (1000 x 12 conditions) were concatenated into a matrix (N x 1000 CT). PCA was then performed on this matrix after z-scoring each row. This bootstrap procedure created a population of trajectories for each condition that captures some information about how much these trajectories may vary. It is worth noting that PCA assumes that the noise in the neurons is uncorrelated and since we are using pseudo-populations, it is possible we are underestimating the correlations between neurons. Noise correlations between neurons can either increase or reduce the separation of these trajectories, which may affect decoding accuracy (Averbeck et al. 2006). To calculate the distance between every pair of trial averaged trajectories for each time point, we computed the euclidean norm between the same time point for every pair of averaged trajectories in the space defined by the top 3 principal components.

#### Geometric Analysis

To understand how the representation of stimulus information in the parietal cortex changes over the course of decision-making, we use the mean-field theoretic manifold analysis technique (Chung et al., 2018; Cohen et al., 2020; Stephenson et al. 2019) to study the geometric properties of the stimulus manifolds, including their manifold capacity, radius and dimensionality. To prepare the data, we counted the number of spikes per neuron in each 80ms time bin for each correct trial for the first 1200ms of each trial. Next, we define 2 object manifolds, corresponding to the 4- and 10-Hz stimulus. We subsampled 50 trials, combining across stimulus durations and animals, for each object manifold. Together, this formed a matrix of size (297 neurons x 2 object manifolds x 50 trials x 14 time points). The neural activations for each stimulus frequency over the 50 trials defines the manifold for that stimulus frequency at each time point. The mean-field theoretic manifold analysis technique then uses this set of activations to compute geometric properties of each object manifold and to evaluate their linear separability. Calculation of each measure was performed using the Replica Mean Field Theory Analysis python library (Stephenson et al. 2019) but we briefly describe the methodology below.

Manifold capacity refers to the maximum number of object manifolds that can be linearly separable given a fixed number of features. If we consider P object manifolds and N neurons, the manifold capacity is defined as *α = P/N.* Intuitively, when *α* is small, then there are few manifolds in a high dimensional space thus making it very easy to find a separating hyperplane for most of the random dichotomy of labels. Alternatively, when *α* is large, there are many object manifolds in a low dimensional space and therefore, it becomes less likely that any dichotomy of manifolds can be linearly separable. The critical manifold capacity, as computed in our analysis, refers to the maximum number of object manifolds P that can be linearly separated given N neurons. This quantity can be estimated from the statistics of anchor points, representative support vectors defining the optimal separating hyperplane, following the methods described in Chung et al. 2018.

In Figure 5, we report manifold capacity relative to its lower bound of 2/M, where M is the number of samples.

Manifold dimensionality, computed from the realized anchor points, estimates the embedding dimension of the manifold contributing to classification. The dimensionality is bounded above by *min*(*M,N*). Since we have M<N, we report manifold dimensionality relative to M in Figure 5D.

Manifold radius is the average distance between the center of the manifold and its anchor points. For linear separability, we care about the size of manifolds relative to how far they are from each other. The manifold radius is therefore reported relative to the norm of the center of the manifold. We also compute the norm of the center of the manifold separately in Figure 5E to estimate how the locations of the manifolds shift over time.

#### Clustering

Neuronal responses were clustered using K-means on a features space comprised of trial-averaged, conditional PSTHs for left-cued and right-cued trials. PSTHs were binned into 10 ms bins, and were then smoothed over a 500ms moving window to reduce noise on the responses (Matlab’s smooth.m function). Each PSTH was then z-scored and combined into a total data matrix, where *N* is the number of neurons and neurons and *T*=151 is the number of data points for each conditional PSTH. PCA was performed on *Z* to reduce the dimensionality of population responses to obtain principal components *W* and score *M*, as. This feature space required *k*=16 components to explain >95% of the covariance in *Z*. The first *k* columns of *M* (data projection onto the top principal components) were used as the feature space for clustering.

The gap statistic criterion was used to determine a principled choice of the best number of clusters (evalclusters.m in Matlab, 5000 samples for reference distribution) (Tibshirani et al., 2001). Specifically, the chosen cluster was defined as the smallest cluster size K, beyond which jumps in gap score Gap(*K*) plateaued and became insignificant,

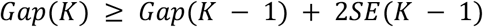

We used the PAIRS statistic to determine if clusters were present in conditional PSTH responses of parietal neurons (Raposo et al., 2014; Hocker et al., 2021). The dimensionality reduced feature space (*i.e.*, the first *k* columns of *M*) were further pre-processed with a whitening transform to yield zero mean and unit covariance. For each data point, the average angle with *n*=4 of its closest neighbors, 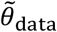, was calculated. This angle distribution was compared with *N*=10,000 sets of independent draws from a reference Gaussian distribution (*N*(0,*I*)), with the same number of data points and the same dimensionality as our data. This *N* datasets were aggregated into a grand distribution, giving the estimated angles *θref*. The number of nearest neighbors *m* is conventionally chosen as the number of neighbors required to give a median nearest neighbor angle *π*/4 for the reference distribution. This number is dependent on the dimensionality of the data, and given the high dimensionality of these responses, even *K*=2 neighbors yielded larger angles than *π*/4. We chose *m* =4 for the results in this work, but note that significant clustering was seen for a wide range of m values (*m*=[2,6]).

PAIRS is a summary statistic of these averages nearest neighbor angles, using the median from the data distribution, 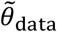 and the median of the grand reference distributions 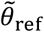:

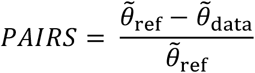

To calculate *p*-values for the PAIRS statistic, reference *PAIRS* values were generated for each of the *N* reference data sets, and the two-sided *p* value (assuming a normal distribution) for the data PAIRS compared to the distribution of reference PAIRS values were reported. We additionally performed a Kolmogorov Smirnov test on the grand reference distribution and data distribution of median nearest neighbor angles.

#### Statistics

Statistical analyses and procedures were implemented with custom-written MATLAB scripts (The Mathworks) that incorporated the MATLAB Statistics Toolbox or in JMP 13.2.0 (SAS). Normally distributed data (as assessed by the Lilliefors test) are reported as mean ± SEM unless otherwise stated. When data were not normally distributed, non-parametric statistical tests were used when appropriate.

## Supplementary Figures

**Supplementary Figure 1.**
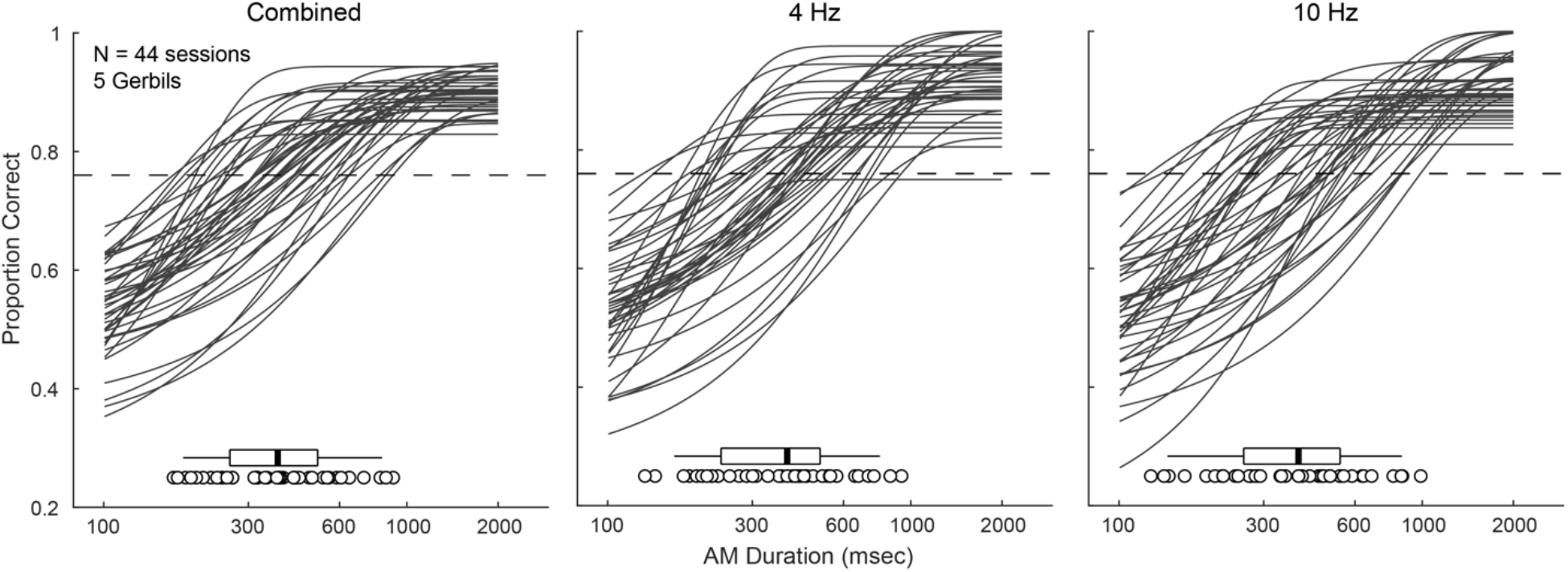
Psychometric functions for all 5 gerbils (44 total sessions) for each AM rate condition. The distribution of minimum integration times for each condition are plotted within each panel. Solid vertical lines within the box and whisker plots represent the median.

**Supplementary Figure 2.**
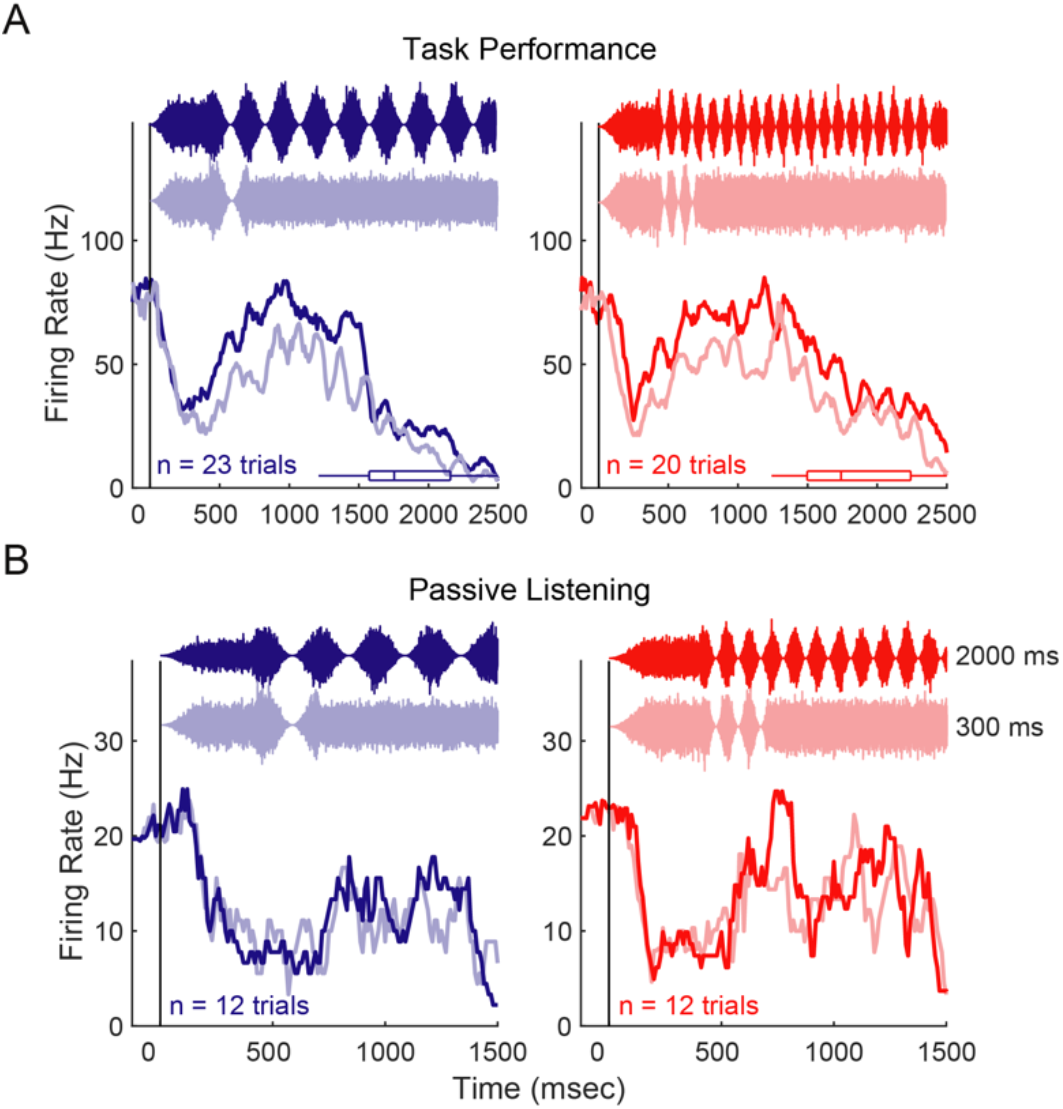
**(A)** Example trial-averaged firing rate post-stimulus time histograms (PSTHs) for one unit recorded during one task performance session across stimulus durations of 300 and 2000 msec. Bin width: 10 msec. Box and whisker plots represent the distribution of response latencies. **(B)** Example trial-averaged firing rate PSTH from the same unit recorded during one passive listening session across stimulus durations of 300 and 2000 msec. Bin width: 10 msec.

**Supplementary Figure 3.**
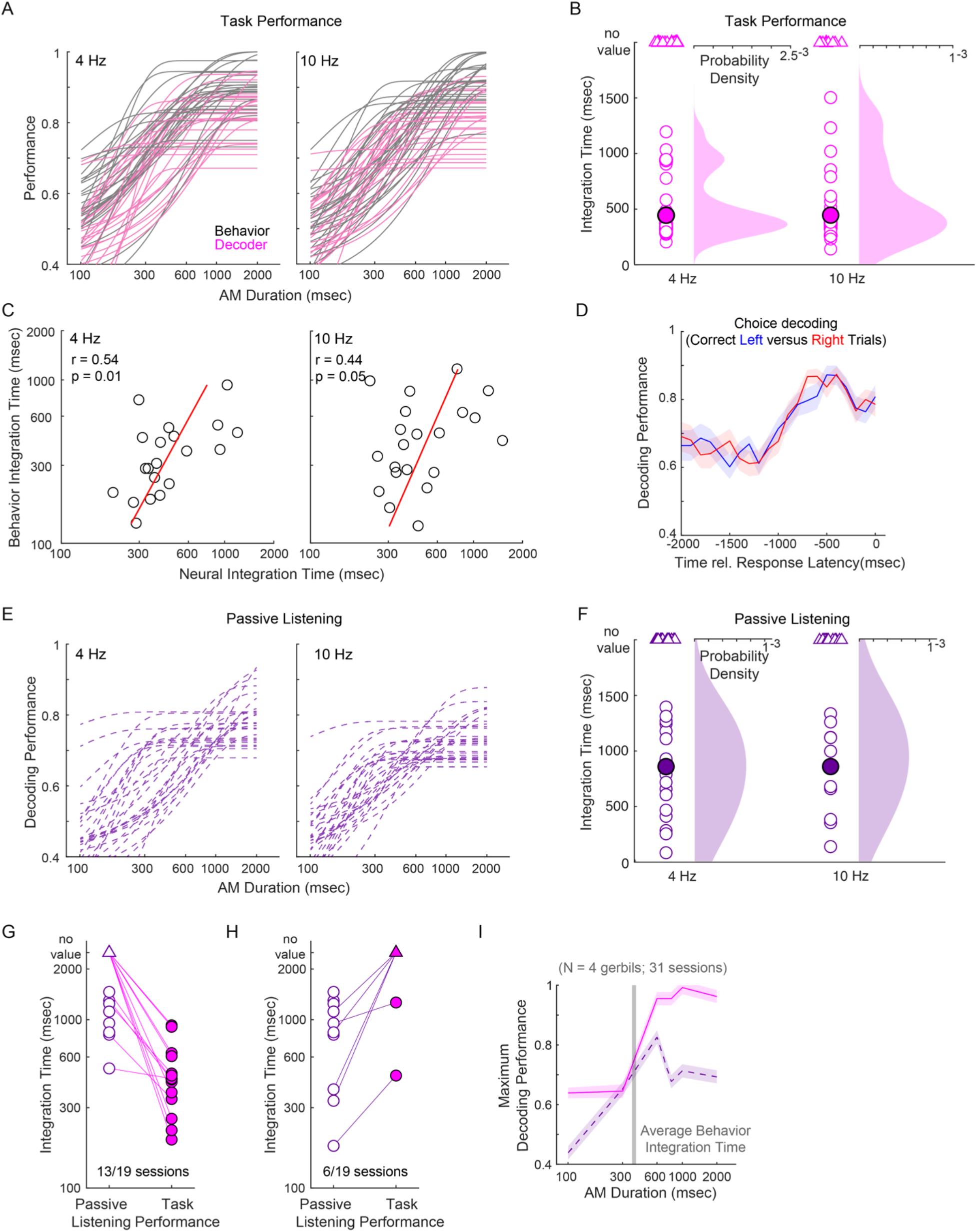
**(A)** Decoding performance and corresponding behavioral performance for each AM rate (4- and 10-Hz) for all sessions. **(B)** Distributions of neural integrations for each AM rate condition. Of the 28 sessions examined, 21 yielded a minimum integration time, while the remaining 7 sessions did not reach the performance criterion on 0.76. There was no significant difference of minimum integration times between acoustic stimuli (Wilcoxon signrank test, p = 0.50; 4-Hz AM median: 402.8 msec; 10-Hz AM median: 407 msec). **(C)** Behavior versus neural integration times for each AM rate condition. Solid red line represents the linear regression. Pearson’s r and statistical significance are noted in the top-left corner of the figure panel. **(D)** Average ± SEM within-session decoding performance for correct Left versus Right trials plotted as a function of time relative to response latency. **(E)** Decoding performance for each AM rate during passive listening for each session. **(F)** Distributions of neural integrations for each AM rate condition during passive listening. Only a portion of passive listening sessions yielded minimum integration times (n = 17/29; maximum decoding performance did not reach 0.76 for the remaining 12 sessions). Of the 17 eligible sessions, there was no significant difference of minimum integration times between acoustic stimuli (Wilcoxon signrank test, p = 0.49; 4-Hz AM median: 829.7 msec; 10-Hz AM median: 838.8 msec). **(G)** Neural integration times between corresponding task performance and passive listening sessions. There were 19 instances where both corresponding task performance and passive listening sessions fit the criterion of 5 simultaneously recorded units. Of those 19 instances, 68.4% (13/19) displayed a decrease in minimum integration times from passive listening to task performance. 8/19 passive listening sessions did not yield a minimum integration time and the remaining 11 sessions produced relatively short minimum integration times (514.5 ± 112.7 msec). **(H)** In 6 cases, integration time diminished or could not be calculated during task performance (passive listening sessions produced minimum integration times of 668.6 ± 217.7 msec). **(I)** Average ± SEM maximum decoding performance across increasing number of units across each stimulus duration for task performance (pink; solid line) and passive listening (purple; dashed line) sessions. Vertical gray bar represents average behavior integration (n = 31 sessions).

**Supplementary Figure 4.**
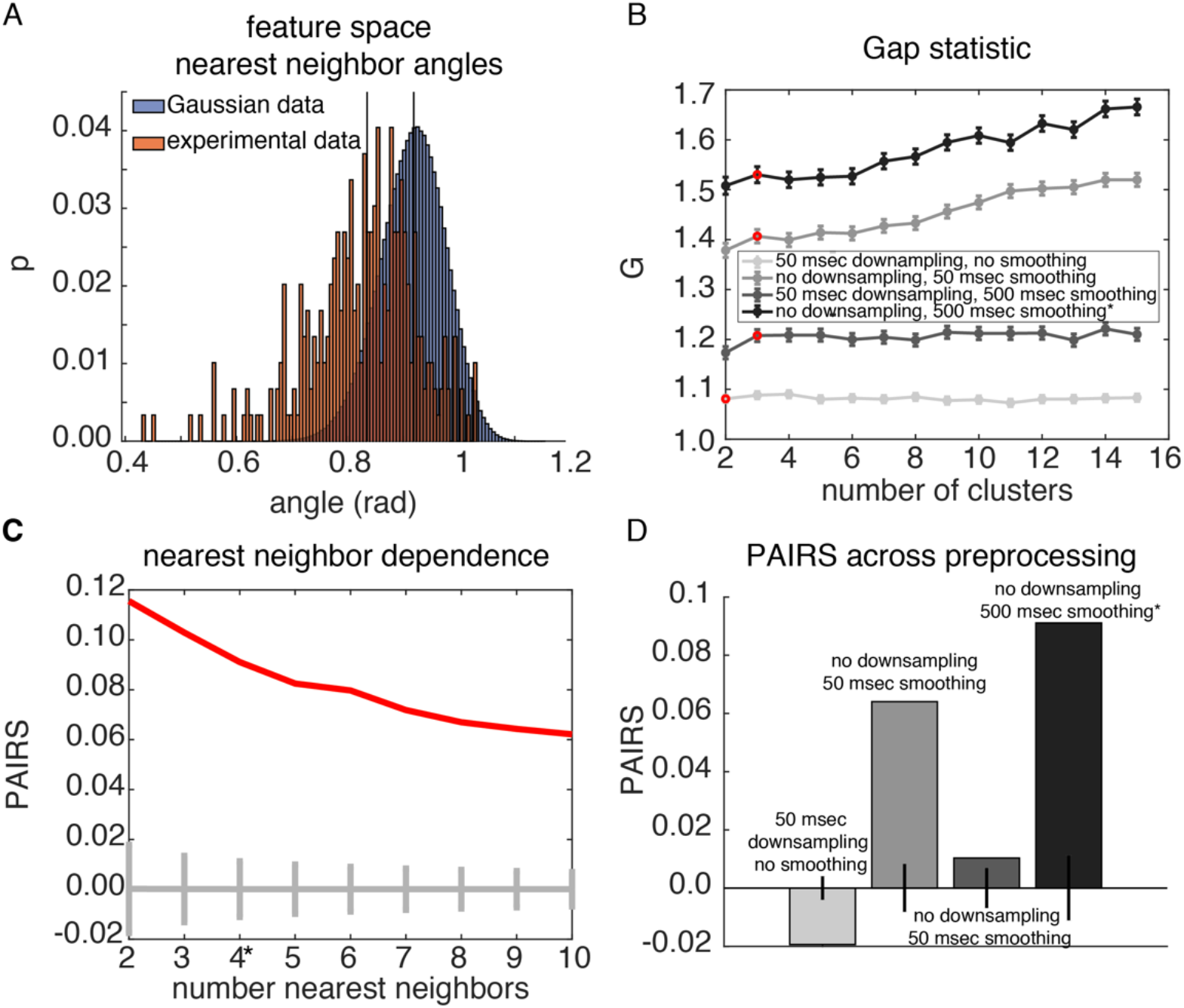
Additional clustering analyses. **(A)** Analysis of angles among nearest neighbors in feature space (red) compared to a null distribution of gaussian data (blue). KS-statistic between these distributions is significantly different (KS test, p = 10^-55^), and PAIRS statistic additionally indicated clustering within the conditional PSTH-based feature space (PAIRS = 0.09, p = 10^-57^). **(B)** Gap statistic test for determining number of clusters across different preprocessing steps of the feature space. Data was z-scored in all cases, with additional downsampling from 10 msec, and/or smoothing over successive time bins. Red dots indicate the number of identified clusters, and error bars denote SEM. 500 msec smoothing without downsampling provided the largest gap statistic values, and this preprocessing was used for clustering in the main figure results. **(C)** PAIRS statistic values for different choices of nearest neighbors size. Red line indicates PAIRS value, and gray line indicates PAIRS value of null distribution, with error bars denoting the 95% confidence interval. 4 nearest neighbors were used to build angle distributions for the PAIRS result in A, though PAIRS values were significant for a large range of numbers of nearest neighbors choice. **(D)** Results of PAIRS test for different preprocessing steps. Error bars denote 95% confidence interval of the null distribution. Only 50 msec downsampling without smoothing indicated a lack of clustering in the data.

**Supplementary Figure 5.**
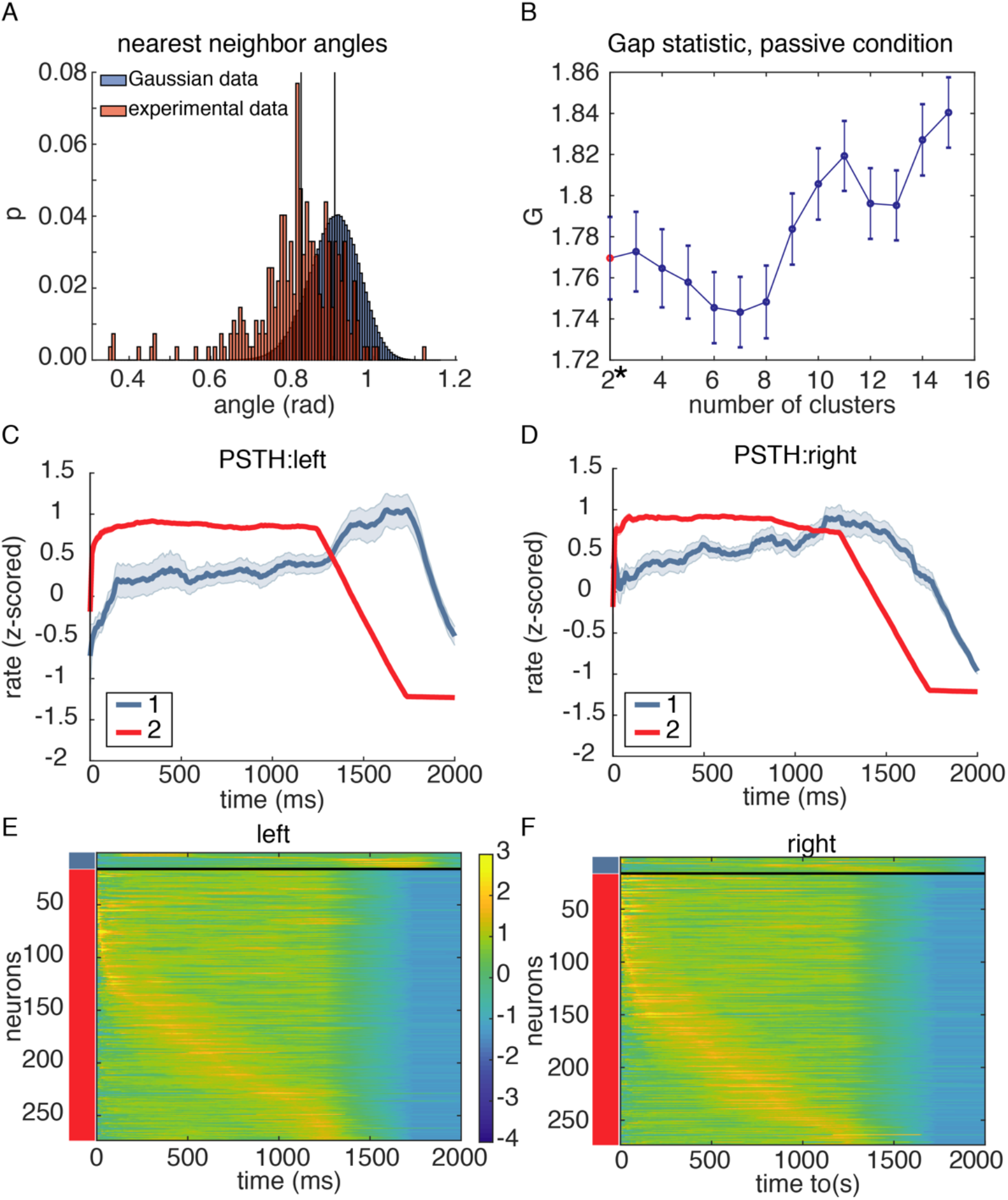
Clustering on conditional PSTHs in the passive listening condition. **(A)** Analysis of angles among nearest neighbors in feature space (red) compared to a null distribution of gaussian data (blue). KS-statistic between these distributions is significantly different (KS-test, p = 10^-49^), and PAIRS statistic additionally indicated clustering within the conditional PSTH (PAIRS = 0.09, p = 10^-47^). **(B)** Gap statistic analysis indicates that two clusters are present in the feature space of conditional responses in the passive condition. **(C-D)** Cluster averaged PSTHs for the 4-Hz stimulus (C) and 10-Hz stimulus (D). **(E-F)** Population PSTHs, grouped by cluster and sorted by time-to-peak within each cluster. Cluster 1 comprises only a small portion of the population. Error bars in all cases are SEM.

## Notes

**Competing Interest Statement**, The authors declare no competing financial interests.

### Competing Interest Statement

The authors have declared no competing interest.

